# Self assembly of HIV-1 Gag protein on lipid membranes generates PI(4,5)P_2_/Cholesterol nanoclusters

**DOI:** 10.1101/081737

**Authors:** Naresh Yandrapalli, Quentin Lubart, Hanumant S. Tanwar, Catherine Picart, Johnson Mak, Delphine Muriaux, Cyril Favard

## Abstract

The self-assembly of HIV-1 Gag polyprotein at the inner leaflet of the cell host plasma membrane is the key orchestrator of virus assembly. The binding between Gag and the plasma membrane is mediated by specific interaction of the Gag matrix domain and the PI(4,5)P_2_ lipid (PIP_2_). It is unknown whether this interaction could lead to local reorganization of the plasma membrane lipids. In this study, using model membranes, we examined the ability of Gag to segregate specific lipids upon self-assembly. We show for the first time that Gag self-assembly is responsible for the formation of PIP_2_ lipid nanoclusters, enriched in cholesterol but not in sphingomyelin. We also show that Gag mainly partition into liquid-disordered domains of these lipid membranes. Our work strongly suggests that, instead of targeting pre-existing plasma membrane lipid domains, Gag is more prone to generate PIP_2_/Cholesterol lipid nanodomains at the inner leaflet of the plasma membrane during early events of virus assembly.

## Introduction

The retroviral Gag protein drives the assembly process of the Human Immunodeficiency Virus type 1(HIV-1) particles [3,5]. This protein is synthesized as a polyprotein Pr55 Gag, which contains three major structural domains, namely matrix (MA), capsid (CA) and nucleocapsid (NC), as well as two spacer peptides, sp1 and sp2 and an unstructured C-terminus p6 peptide. Each of these domains are known to have a specific and distinct function during the viral assembly process. Importantly, the N-terminal MA domain targets Gag to the plasma membrane and mediates membrane binding, the CA domain is responsible for Gag-Gag interaction and self-assembly and the NC domain recruits the viral RNA that also acts as a scaffold for viral particle assembly [42]. Although it has recently been shown that HIV assembly may be initiated in the cytosol [28], it is commonly accepted that the formation of large HIV-1 assembly complex mainly occurs at the plasma membrane (PM) of the virus producing cells. Two main features of the N-terminal matrix (MA) domain of Gag govern the HIV-1 Gag inner leaflet PM binding: the N-terminal myristate and the highly basic region (HBR) that contains a specific binding pocket for the phosphatidyl inositol (4,5) bisphosphate lipid (PI(4,5)P_2_ or PIP_2_) [48]. PIP_2_ has been shown to play a major role for HIV-1 assembly in cells since cellular depletion of PIP_2_ decrease the efficiency of viral particle release [43]. In addition, PIP_2_ is also enriched in the virus envelope relative to the PM of the virus-producing cells [10]. This lipidomic analysis also shows that HIV particles are enriched in sphingolipids and cholesterol [10] supporting the idea that the viral particles are released from the so called “rafts” domains. Historically, Gag has been shown to associate with detergent resistant membrane (DRM) by cell membrane fraction assays [44], [7]. Nevertheless, the exact composition of these “rafts” or DRM is controversial [8,29]. Indeed, many different types of rafts may exist within the plasma membrane as long as they are enriched in cholesterol and sphingolipids [45]. However, Gag self-assembly occurs at the inner leaflet of the cellular PM where sphingolipids are poorly present. Another concern is the ability of PIP_2_ to partition into rafts while natural PIP_2_ has an polyunsaturated fatty acyl residue at the sn-2 position of the glycerol backbone. Based on NMR data using truncated acyl chains PIP_2_, Saad *et al.* suggested a model to circumvent such constrain [48]. In this model, the sn-2 acyl chain of the PIP_2_ is removed from the plane of the membrane and trapped into an hydrophobic pocket of MA. However, coarse grained dynamics studies [11] and new NMR experiments using full length acyl chain PIP_2_ [40] showed the opposite. Moreover, recent experiments using either a multimerizable matrix domain of HIV-1 Gag [34] or RSV Gag [54] also exhibited contradictory results regarding their partitioning in lipid domains of giant unilamellar vesicles (GUVs). Therefore, the ability of the Gag/PIP_2_ complex to partition preferentially into “raft” domains or more generally into PM pre-existing domains to enhance virus assembly is still a matter of controversy [36]. In order to elucidate whether Gag would bind to pre-existing “rafts” or lipid domains in the PM or generate its own lipid domains for assembling, we first decided to monitor its binding and partitioning to single or dual-phase model membranes made with simple and complex lipid compositions (see table1 for detailed composition of lipid mixtures). On GUVs, we observed that Gag was mainly partitioning into *L_d_* phase, i.e. more likely out of “rafts” microdomains in the dual-phase model membrane and that Gag was not generating micrometer range lipid phase separation in single phase model membranes. We therefore monitored, at the nanoscale level, a possible PIP_2_, cholesterol (Chol) and sphingomyelin (SPM) reorganization during Gag self-assembly. Unfortunately, direct imaging of lipid nanodomains generation is a not an easy task. Even the presence of pre-existing lipid nanodomains has indirectly been revealed by monitoring lipid diffusion [2, 17, 20, 53, 55]. Although accurate in detecting and measuring the size of pre-existing nanodomains, these methods are not fast enough to follow the generation of nanodomains. Therefore, we decided to use indirect approaches such as fluorescence quenching [22,46] or Forster Resonant Energy Transfer (FRET) [27]. FRET or self-quenching occurs when two (similar or different) fluorescent molecules are found together within a 3 to 10 nm radius. Starting from a simple lipid mixture and using self-quenching experiments of respectively PIP_2_, Chol and SPM, we monitored the generation of Gag induced lipid nanodomains. We then performed the same type of experiments with dual labelling of the lipids (Chol and PIP_2_, or SPM and PIP_2_) and complete them with FRET experiments to analyse how Gag was differently sorting these lipids during its self-assembly. We finally extended this study to more complex lipid compositions mimicking either lipid “rafts” or inner leaflet of the PM. Our results show that Gag self-assembly is able to generate PIP_2_ nanodomains on model membranes whatever was the composition we tested here. Importantly, these nanodomains contain Chol but not SPM.

## Results

Since PIP_2_, Chol and SPM have been shown to be enriched in the virus envelope [10], the impact of Gag self-assembly on their lateral distribution was tested in different types of model membranes (Fig.1A) exhibiting different lipid compositions. Large unilamellar vesicles (LUVs) were firstly used in order to easily control the protein over lipid molecular ratio. Nevertheless, because HIV-1 Gag is expected to assemble at the plasma membrane, we thereafter mainly used supported lipid bilayers (SLBs)as membrane models. Gag is known to bind specifically PIP_2_ [13]. We then started with a simple PIP_2_ containing lipid mixture (PC/PS/PIP_2_ hereafter called “basic”) in which we introduced Chol or SPM (hereafter called “substituted basic”). Obviously, cellular PM have more complex lipid compositions. We therefore decided to extend our study to compositions mimicking either the inner leaflet cellular plasma membrane [32] (hereafter called “inner leaflet”) or the “lipid rafts” domains (i.e. two separated phases, a liquid disordered (L*_d_*) and a liquid ordered (L*_o_*) enriched in cholesterol and sphingomyelin) [34] (hereafter called “rafts-mimicking”). Table 1 gives the exact compositions of the lipid mixtures used in this study.

**Figure 1.**
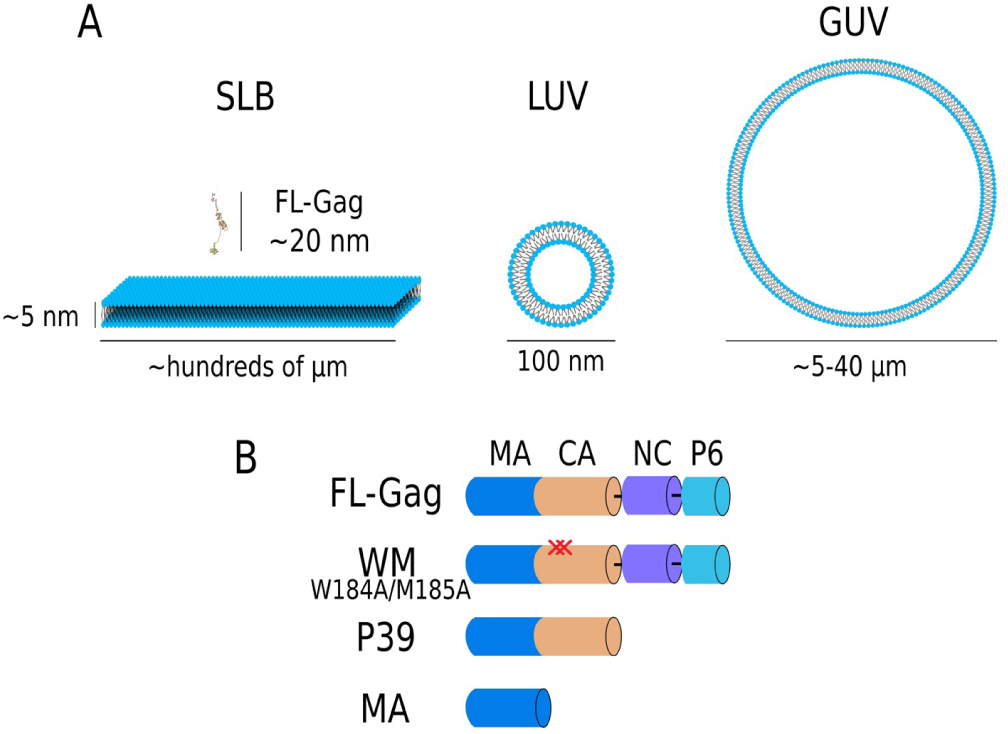
Scheme of the different lipid membranes and proteins used in this study. **A**: Representation of the model lipid membranes and of an elongated FL-Gag protein used in this study (see Table 1 and SI Table S1 for all the different lipid compositions used in this study). **B**: Schematic representation of the different Gag mutants used in this study, emphasizing the mutations and the differences in the domains present for each mutant

**Table 1.**
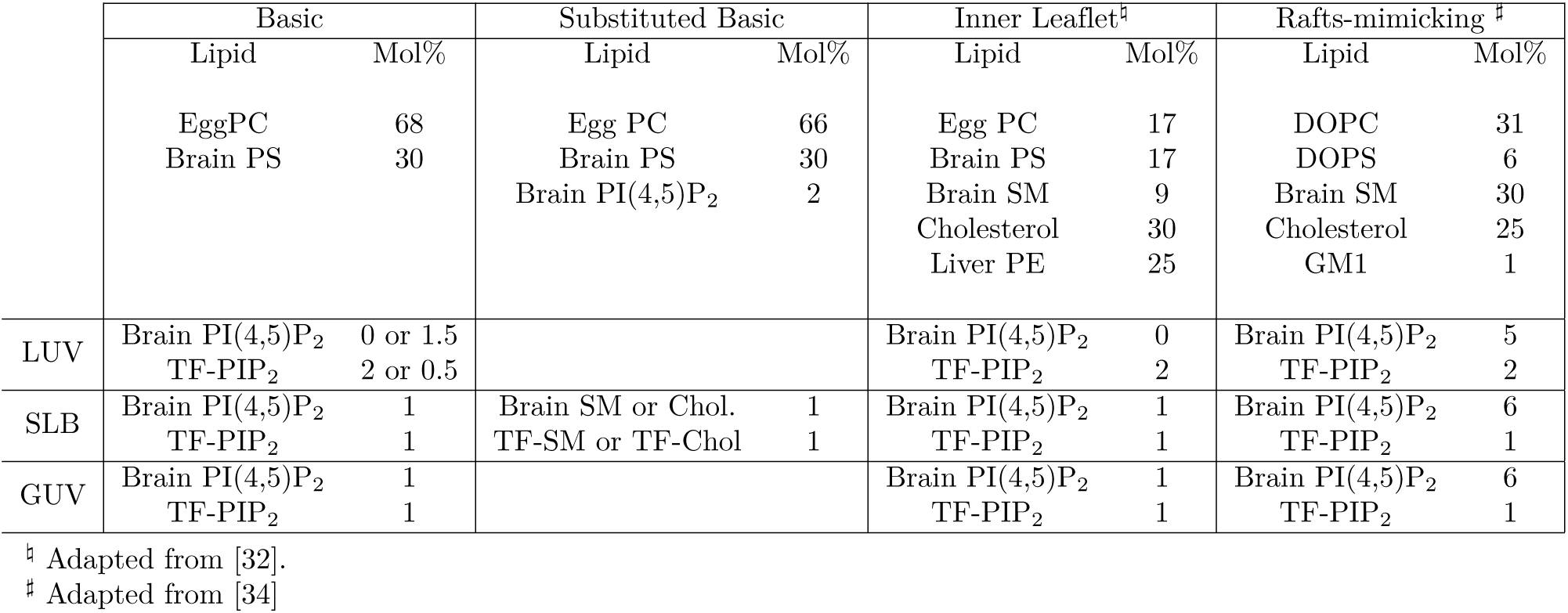
Different lipid composition of LUV, SLB and GUV used in the single labelled lipid experiments

To assess a potential reorganisation of these lipids during Gag self-assembly, we used full-length Gag (FL-Gag) and different mutants. Theses mutants include: (1) a mutant of CA involved in CA-CA interaction and Gag self-assembly (WM) [15](2) Gag lacking the C-terminus NC-sp2-p6 domains that is involved in NC-RNA association (P39). The membrane binding domain alone of Gag (MA) was also tested (see Figure1B). Finally, a cellular PM PIP_2_ binding protein, the PH domain of EFA6 (PH-EFA6) [38] and a peptide, MARCKS (151-175) that is known to laterally redistribute PIP_2_ [22] were used as controls.

### HIV-1 Gag efficiently binds model membranes containing PIP_2_

The ability of the different proteins and peptide to bind LUVs and SLBs was analysed using membranes of “basic” composition. For this purpose, we monitored protein binding with two different techniques. On one hand, we performed LUV cosedimentation assays at fixed protein concentration (1 *µ*M) and increasing PIP_2_ concentrations. After 15 min of incubation, LUV bound proteins were separated from unbound proteins by ultra-centrifugation. The pellet (P, bound proteins) and the supernatant were deposited on a denaturating gel to quantify the amount of bound and unbound proteins (Figure 2A). On the other hand, we used Quartz Crystal Microbalance (QCM) to monitor increasing protein concentrations (from 10^−2^ to 10 *µ*M) association to SLBs at fixed PIP_2_ concentration. SLBs was formed onto the quartz crystal. Proteins were then injected and the change in the resonance frequency of the quartz crystal was monitored (Figure 2B). This change in frequency is directly connected to the mass increase at the surface of the quartz crystal (see, for details).

**Figure 2.**
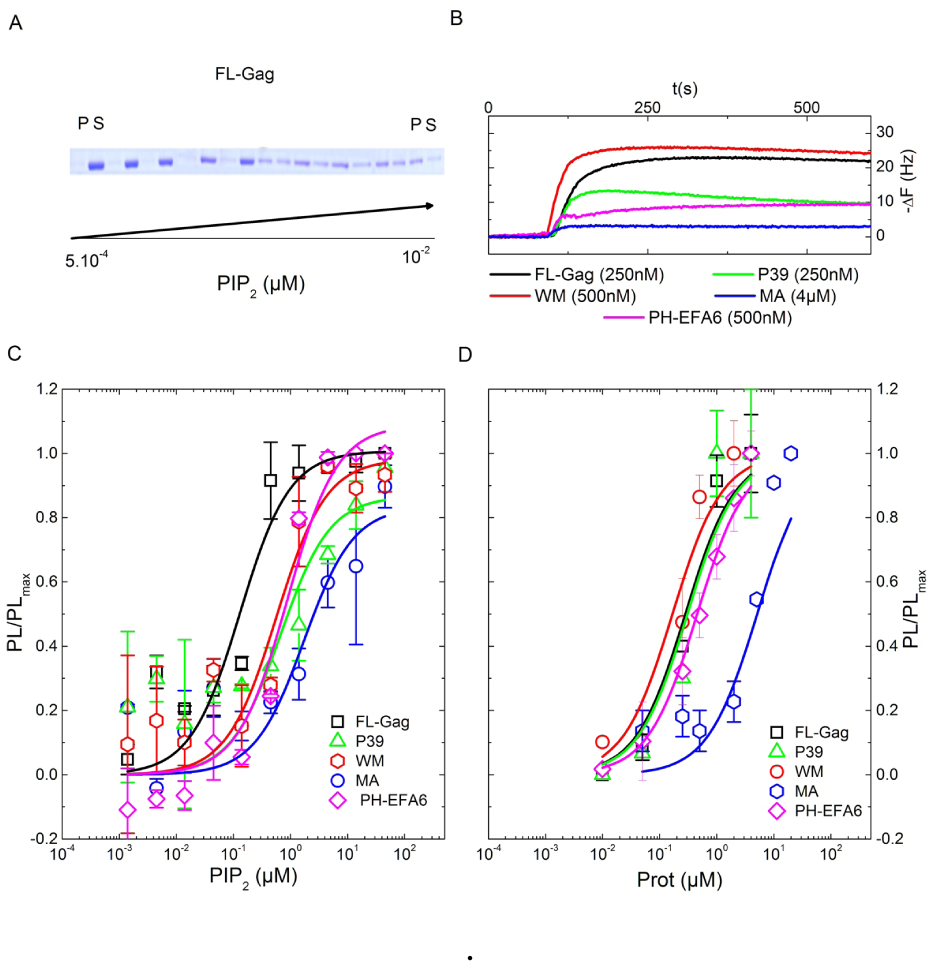
Binding of FL-Gag, its mutants and PH-EFA6 to “basic” lipid mem-b ranes. **A**: Typical SDS-PAGE obtained after co-sedimentation assay for FL-Gag with increasing concentrations of LUV containing 2% mol PIP_2_ (P:Pellet, S:Supernatant). **B**: Typical change in resonance frequency observed during QCM-D experiments after addition of the different proteins at different concentrations on a basic composition SLB. **C and D**: Binding isotherm curve obtained from co-sedimentation assays (**C**, n=3) and QCM-D experiments (**D**, n=2) for the different proteins used in this study. Experimental values were fitted using equation 1. K_p_ obtained from these binding isotherms are summarized in table 1

The ratio of membrane bound to unbound protein is linked to an apparent partition coefficients (*K_p_). K_p_* were determined by fitting with eq.1 the plots of membrane bound protein with either increasing concentrations of lipids (Figure 2C) or increasing concentration of protein (Figure 2D).

Except for MA, *K_p_* obtained for other proteins were less than 1*µM*. The results, summarized in table 2, are in good agreement with the previously obtained *K_p_* values for some of the different proteins used in this study. Interestingly, the *K_p_* values of MARCKS and PH-EFA6 (0.7 < *K_p_* < 0.8*µM*) were found to be in the range of the one obtained for Gag (FL) and its mutants (0.2 < *K_p_* < 0.8*µM*). This shows that the ratio of membrane bound over total protein will be equivalent for all these proteins, allowing therefore direct comparisons of their respective role on the lateral sorting of PIP_2_ after membrane binding.

**Table 2.** Apparent K_p_ of the different proteins used in this study

| Protein | $K_p(\mu M)$<br>LUV | $K_p(\mu M)$<br>SLB | $K_p(\mu M)$ | Lip. Comp (mol %) | Ref. |
| --- | --- | --- | --- | --- | --- |
| FL | $0.13 \pm 0.05$ | $0.40 \pm 0.11$ | $0.88 \pm 0.20$ (LUV) | POPS (100) | [18] |
| P39 | $0.66 \pm 0.44$ | $0.47 \pm 0.26$ | | | |
| WM | $0.56 \pm 0.26$ | $0.26 \pm 0.13$ | | | |
| MA | $1.72 \pm 1.05$ | | $2.46 \pm 1$ (LUV) | POPS (100) | [18] |
| | | $7.16 \pm 3.33$ | $8.2 \pm 0.7$ (SLB) | DOPC:DOPS:PI(4,5)P<sub>2</sub> (80:15:5) | [4] |
| PH-EFA6 | $0.88 \pm 0.27$ | $0.68 \pm 0.03$ | 0.5 (SLB) | PC:PE:PS:PI(4,5)P<sub>2</sub><br>(34:34:30:2) | [38] |
| MARCKS | n.d.<sup>#</sup> | $0.65 \pm 0.23$ | $4.4 \pm 1$ (LUV) | PC:PI(4,5)P<sub>2</sub><br>(99:1) | [52] |

<sup>#</sup> Could not be determined by this method.

### FL-Gag is mainly partitioning to L*_d_* phases and does not induce micrometer phase separation

We firstly monitored the possible existence of Gag induced micrometer-range PIP_2_ domains by imaging the surface distribution of a fluorescently labelled PIP_2_ (namely Top-Fluor PIP_2_ or TF-PIP_2_) and a fluorescently labeled FL-Gag (FL-Gag Alexa 549 or A549-FL-Gag) in “basic” GUVs. As expected, both TF-PIP_2_ and A549-FL-Gag showed a uniform distribution over the surface of the GUV (fig.3A). The same uniform distribution was also observed with “inner leaflet” GUVs (fig. 3A). Spatial auto-correlation of the fluorescence of either PIP_2_ or FL-Gag (fig.3B) both exhibited fast decorrelation showing that the fluorescent labelling of PIP_2_ is not inducing micrometer partitioning *per se* and that, if they exist, the PIP_2_ clusters are less than 200 nm size (i.e. below the diffraction limit). Interestingly, labelled FL-Gag was essentially restricted to L*_d_* phases in the “rafts-mimicking” GUVs (figure 3C and D), confirming what was already observed by Keller *et al.* [34] with a multimerized derivative of MA. This strongly suggested that FL-Gag preferentially binds and remains in non “rafts” lipid phases.

**Figure 3.**
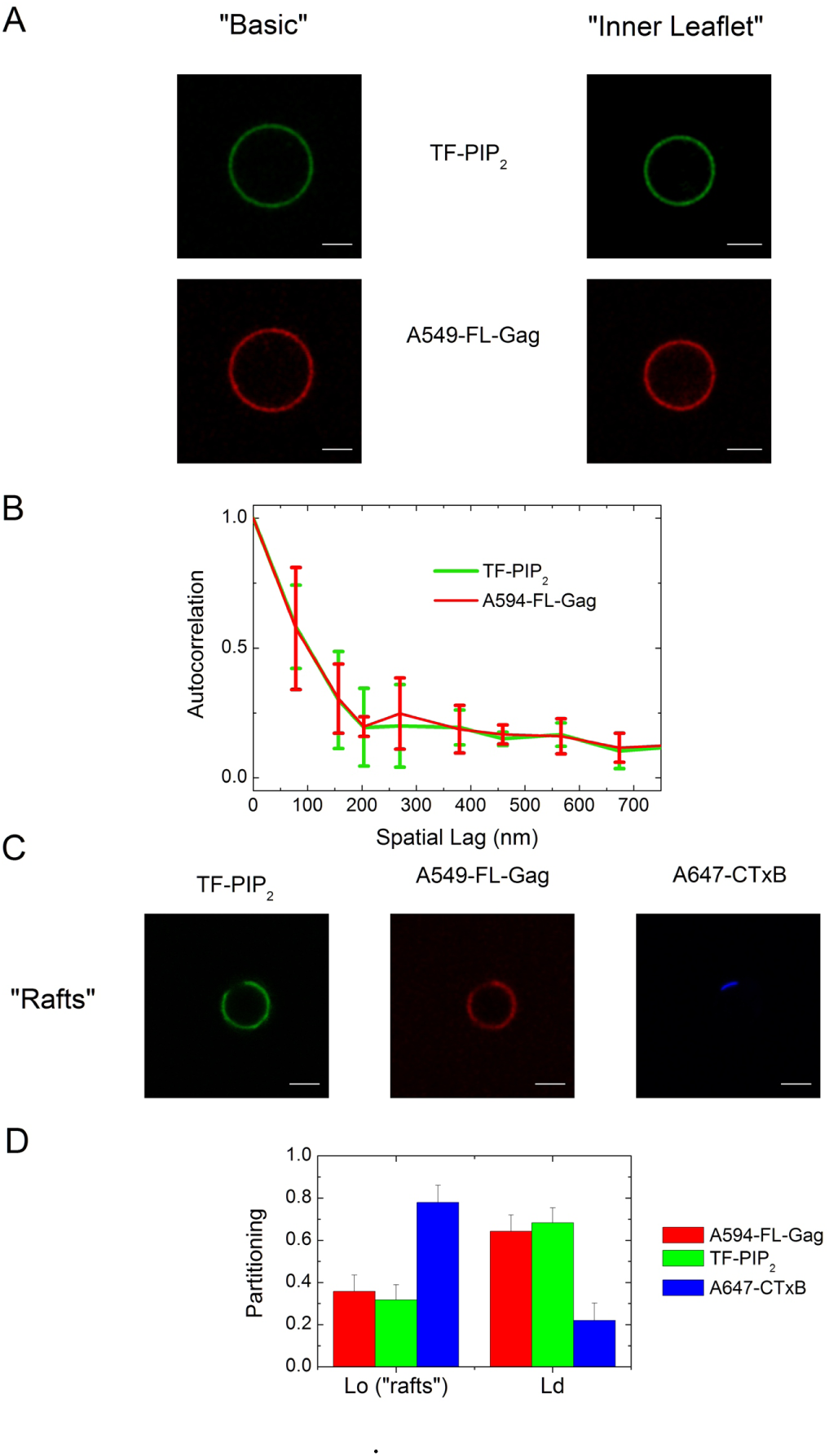
Fluorescence distribution of TF-PIP_2_ and A594-FL-Gag on “Basic”, “Inner Leaflet” and “rafts-mimicking” GUVs. **A**: Left, localization of TF-PIP_2_(green) and Alexa 594 FL-Gag(A594-FL-Gag)(red) in a “Basic” composition GUV. Right, localization of TF-PIP_2_(green) and Alexa 594 FL-Gag(A594-FL-Gag)(red) in an “Inner Leaflet” composition GUV, (scale bar 5 µm) **B**. Spatial autocorrelation of the fluorescence intensities of TF-PIP_2_(in green) and A594-FL-Gag(in red) for the “Basic” composition calculated using eq. 2, (mean±s.d., n=4). **C**: Localization in “Raft” GUVs (**F**) of TF-PIP_2_(in green), A594-FL-Gag (in red) and GM1, a raft partitioning gan-glioside, labeled with alexa 647 cholera toxin B (A647 Ctx-B)(in blue), (scale bar 5 µm). **D**: Partitioning in L_o_ and L_d_ phase of “raft” GUVs for A-594 FL, TF-PIP_2_ and A647-CtxB (mean ± s.d., n=25) as determined using eq.3

### HIV-1 Gag self-assembly induces PIP_2_ nanoclustering

The lack of micrometer scale PIP_2_ induced domain lead us to monitor, using self-quenching experiments, a possible PIP_2_ lateral redistribution upon Gag self-assemby on LUVs or SLBs. As stated in the introduction, self-quenching occurs when two fluorescent molecules are close to each other (5*<*d*<*10 nm). To perform these experiments, we first controlled that increasing concentrations of MARCKS induced an increasing quenching of the TF-PIP_2_ as it was reported before with Bodipy TMR-PIP_2_ (BT-PIP_2_) [22] (See SI Fig.S1).We then monitored the change in TF-PIP_2_ fluorescence after addition of Gag or its mutant on “basic” LUVs in order to correctly control the protein to accessible PIP_2_ ratio 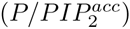. We observed an increase in TF-PIP_2_ fluorescence (fig.4A) opposite to MARCKS. From figure 4B it was evidenced that addition of increasing concentrations of FL-Gag, P39 and WM induced an increasing fluorescence unquenching of TF-PIP_2_. On the other hand, adding increasing concentrations of MA did not generate any change in the fluorescence of TF-PIP_2_. Identically, PH-EFA6, known to specifically bind PIP_2_ without reorganizing its lateral distribution, did not produce any effect either (fig.4A and B). A major difference between the Gag matured protein MA and the precursor proteins (FL-Gag, P39 and WM) is their capacity to form large oligomer complexes via the CA-CA interactions [41], although WM is less efficient (two orders of magnitudes in solution) in supporting the formation of large Gag lattice due to its inability to dimerize with nearby CA hexamer [15]. Interestingly, this fluorescence unquenching was always less efficient in the case of WM compared to FL-Gag (from 3 to 5 times), indicating that this fluorescence variation depended on the capacity of Gag to multimerize.

**Figure 4.**
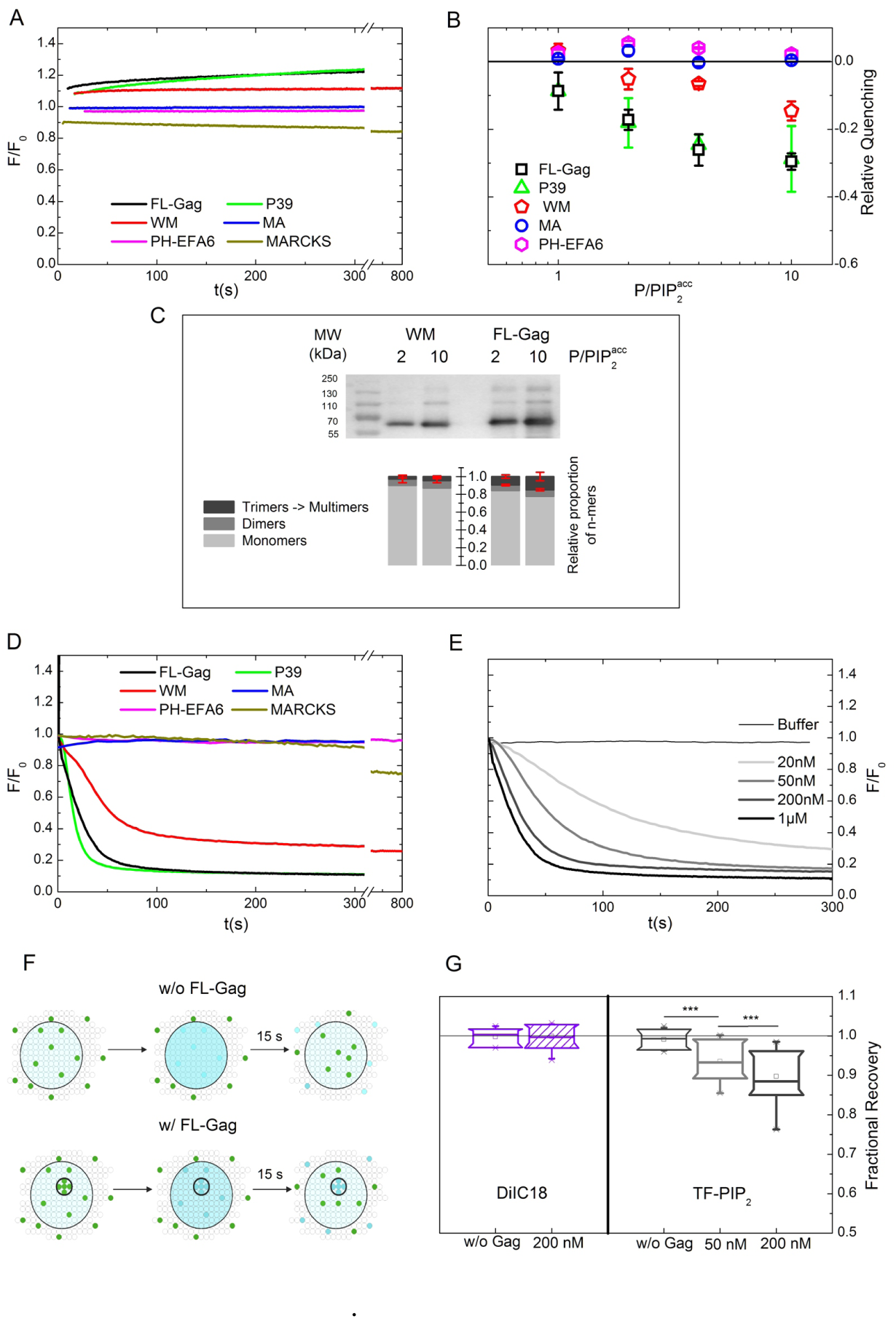
PIP_2_ nanoclusterization induced by Gag self-assembly on basic composition lipid membranes. **A**: Typical time course of TF-PIP_2_ fluorescence on LUVs after addition of the different proteins or peptide at a 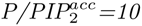. **B**: Relative quenching values observed for FL-Gag, its mutant, MA and PH-EFA6 on LUVs (mean ± s.d. values of n ≥ 3 for each 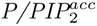 conditions, except P39 (2 ≤ n ≤ 3)). **C**: Differences in WM and FL-Gag self-assembly efficiency on LUV at two different 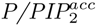 ratio. Bar graph represents the mean signal observed for monomers dimers and trimers & multimers obtained from independent experiments (mean ± s.d., n=3). **D**: Typical time course of TF-PIP_2_ fluorescence on SLBs after addition of 1 µM of the different proteins or peptide. **E**: Fluorescence time course of TF-PIP_2_ after addition of increasing FL-Gag concentrations. **F**: Schematic representation of the effect of an immobile fraction on the fractional recovery. Upper part, without Gag, fractional recovery = 1. Lower part, with Gag, self assembled Gag trapped PIP_2_ are still in the bleached area leading to a fractional recovery < 1. **G**: Plot box of the fractional recoveries obtained from FRAP measurements before and after addition of increasing concentrations of FL-Gag on SLB containing either DiIC18 as a lipid analogue control (left) or TF-PIP_2_ (right).(Boxes are 25,75% with bars max and min values of n ≥ 15, ***: p ≤ 10^−3^ for Student t-test at 0.01 confidence level)

To support this hypothesis, FL-Gag and WM were incubated with “basic” LUVS for 15 minutes followed by ultracentrifugation. The self-assembly states of WM and FL-Gag bound to LUVs were analysed by performing a non denaturating gel electrophoresis to preserve existing multimers, followed by immunoblot. Figure 4C showed that, in the two 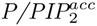 ratio tested, WM was at least two times less efficient than FL-Gag in multimerizing on membranes (fig.4B). This ratio was in the range of the relative efficiency of TF-PIP_2_ unquenching observed between FL-Gag and WM. This confirms the role of Gag self-assembly in the observed TF-PIP_2_ fluorescence changes. Although LUVs provide a simple, reliable and easy to control (in terms of protein to lipid ratios) model membranes, Gag self-assembly usually occurs on membranes with higher radius of curvature. Therefore, time course fluorescence self-quenching of TF-PIP_2_ were also conducted on “basic” SLBs. For FL-Gag and P39 the TF-PIP_2_ fluorescence intensity changed on SLBs (fig. 4D) and was opposite to the one on LUVs (fig. 4A) while still being a function of the protein concentration (figure 4E for FL-Gag). As it was the case for LUV, the TF-PIP_2_ quenching induced by addition of the WM mutant was less efficient than in the case of FL-Gag. PH-EFA6 and MA did not induced changes in TF-PIP_2_ fluorescence (fig. 4D). Interestingly 4D shows that MARCKS addition induced the same TF-PIP_2_ quenching on SLBs and LUVs (see fig.4A), suggesting that the differences observed with Gag and its mutant depend on model membrane curvature (See SI fig. S2 for detailed explanation). Altogether these results show that the self-assembly of HIV-1 Gag is generating TF-PIP_2_ clusters.

Diffusion of Gag is expected to decrease upon self-assembly. Hendrix et al. [28] already observed that Gag multimers exhibited a diffusion coefficient(D) of 0.01 *µm*^2^.*s*^−1^ at maximum. We controlled that this decrease in diffusion was also observed for TF-PIP_2_. For that purpose, we performed Fluorescence Recovery After Photobleaching (FRAP) experiments before and after addition of FL-Gag on the same SLBs that the one used for the quenching experiments. Considering that PIP_2_ clustered in Gag multimers will diffuse with the same coefficient than Gag multimers, then, the fluorescence of PIP_2_ in a 1*µ*m radius bleached area containing multimers should not totally recover after 15s (fig.4F). This will lead the fractional fluorescence recovery value (defined in eq.4) to be less than 1. We therefore monitored the change in fractional fluorescence recovery in the absence or presence of FL-Gag. In the absence of FL-Gag, fractional fluorescence recovered to 1, as expected for freely diffusing lipids (fig.4G, right). On the opposite, addition of increasing FL-Gag concentrations induced a decrease in PIP_2_ fluorescence fractional recovery as well as an increase in fluorescence quenching (previously shown in fig.4E). We controlled that the decrease in fractional recovery was specific of PIP_2_. Indeed, we did not observe any change in the fractional fluorescence recovery of a lipid analogue fluorescent dye (diIC18, see materials in SI) in the presence or in the absence of Gag (fig.4G, left).

Our data clearly show that HIV-1 Gag is sorting PIP_2_ in the lipid membrane and that Gag self-assembly generates PIP_2_ nanoclusters in model membranes.

### Cholesterol but not sphingomyelin is sensitive to HIV-1 Gag self-assembly

Since Chol and SPM are the main lipid components of raft domains, their enrichment into the Gag self-assembly induced PIP_2_ nanoclusters was also assessed. 2% mol of egg phosphatidyl choline (EPC) present in our “basic” lipid composition, were substituted either by SPM or by Chol with half of them being labelled with TF derivatives (see SI Table1 for detailed composition). This substitution allowed the net surface charge and the PIP_2_ content to be maintained, limiting any drastic change in the partitioning constant *K_p_*. Time course fluorescence change of either TF-SPM or TF-Chol were then monitored after FL-Gag addition.

Figure 5A shows that FL-Gag addition had no effect on TF-SPM fluorescence whereas figure 5B shows that increasing concentrations of FL-Gag induced increasing quenching of TF-Chol, as it was the case for TF-PIP_2_ (Fig.4E). This shows that FL-Gag self- assembly is able to generate Chol-enriched lipid nanodomains whereas it is not changing the SPM lateral distribution.

**Figure 5.**
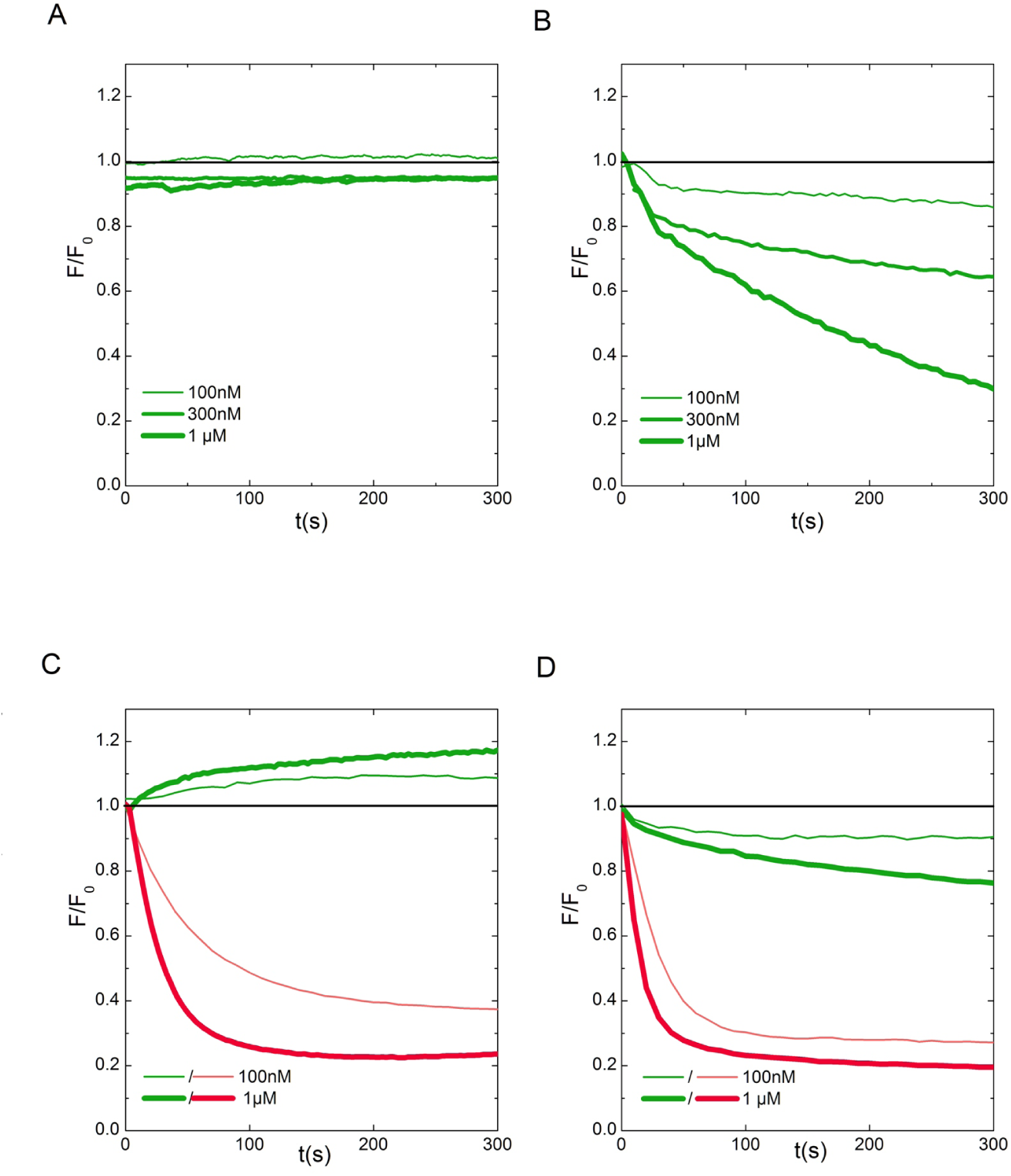
Gag self-assembly is sorting cholesterol but not sphingomyelin. **A & B**: Typical fluorescence time course after addition of increasing FL-Gag concentrations on sub-situted basic SLBs labeled with TF-SPM (**A**) or TF-Chol (**B**). **C & D**: Simultaneous fluorescence time course after addition of increasing concentrations of FL-Gag on substituted basic SLBs labelled with TF-SPM (in green) and BT-PIP_2_ (in red) (**C**) or TF-Chol (in green) and BT-PIP_2_ (in red) (**D**)

Co-clustering of these molecules was also assessed by simultaneously monitoring the change in fluorescence upon addition of FL-Gag with either TF-SPM and BT-PIP_2_ (fig.5C) or TF-Chol and BT-PIP_2_ (fig.5D) labelled SLBs (see SI table 1 for exact dual labelled lipid membranes compositions).

TF-Chol and BT-PIP_2_ exhibited simultaneous quenching after addition of various concentration of FL-Gag, suggesting that TF-Chol and BT-PIP_2_ are co-clustered in Gag self-assembly induced nanodomains. While BT-PIP_2_ was quenched as previously, TF-SPM exhibited a surprising apparent fluorescence unquenching after Gag addition. This was an unexpected result regarding the TF-SPM quenching induced by FL-Gag self-assembly in the presence of unlabelled PIP_2_ (fig. 5A). Since this apparent unquenching of TF-SPM upon addition of FL-Gag only occurred when BT-PIP_2_ was present (see fig 5A vs fig 5C), we checked if TF-SPM (as a donor) and BT-PIP_2_ (as an acceptor) were able to exhibit FRET. Donor FRET efficiency has been shown to be a function of acceptor concentration in a lipid bilayer [21]. This means that the FRET efficiency will decrease if acceptor (BT-PIP_2_) local concentration decreases and that, as a consequence, the fluorescence of the donor (TF-SPM) will increase. We ensured this was the case by performing acceptor photobleaching experiments in order to decrease the acceptor concentration, locally and reversibly. We observed that, in the area where the acceptor (BT-PIP_2_) was bleached, the fluorescence of the donor (TF-SPM) increased. This effect was abolished when concentrations were re-equilibrated by diffusion (see SI Fig3). In our TF-SPM/BT-PIP_2_ co-clustering assay, we observed a decrease in BT-PIP_2_ due to Gag self-assembly induced quenching and concomitantly, a non reversible TF-SPM fluorescence increase due to loss in FRET efficiency induced by decreasing concentration of acceptor (TF-PIP_2_), showing that Gag self-assembly clusters BT-PIP_2_ without TF-SPM.

Altogether, these results show that HIV-1 Gag self-assembly is generating PIP_2_ and cholesterol nanodomains while excluding sphingomyelin.

### HIV-1 Gag is driving PIP_2_ and cholesterol nanoclustering independently of surrounding lipids

Because cellular PMs are complex lipid mixtures, we examined a possible role for this complexity on the lateral sorting of PIP_2_, Chol and SPM during HIV-1 Gag self-assembly. For that purpose, we used the two other different lipid mixture mimicking either the inner leaflet of cells PM (”inner-leaflet”) or the “rafts” lipid mixture (”raft-mimicking”)(see Table1 and SI Table 1 for detailed lipid composition).

We first compared the effect of FL-Gag self-assembly on PIP_2_ clustering for these three lipid compositions. No characteristic change in TF-PIP_2_ fluorescence unquenching with increasing concentrations of FL-Gag (LUVs, fig.6A) or fluorescence quenching (SLBs, fig.6B) could be clearly detected amongst the three different lipid compositions. We then tested again Chol and SPM ability to partition into these FL-Gag induced PIP_2_ nanoclusters in the complex lipid mixtures. Figure 6C (TF-SPM/BT-PIP_2_) and 6D (TF-Chol/BT-PIP_2_) exhibited the same tendency for the fluorescence time courses of labelled lipid upon FL-Gag addition on SLBs, independently of the lipid composition.

**Figure 6.**
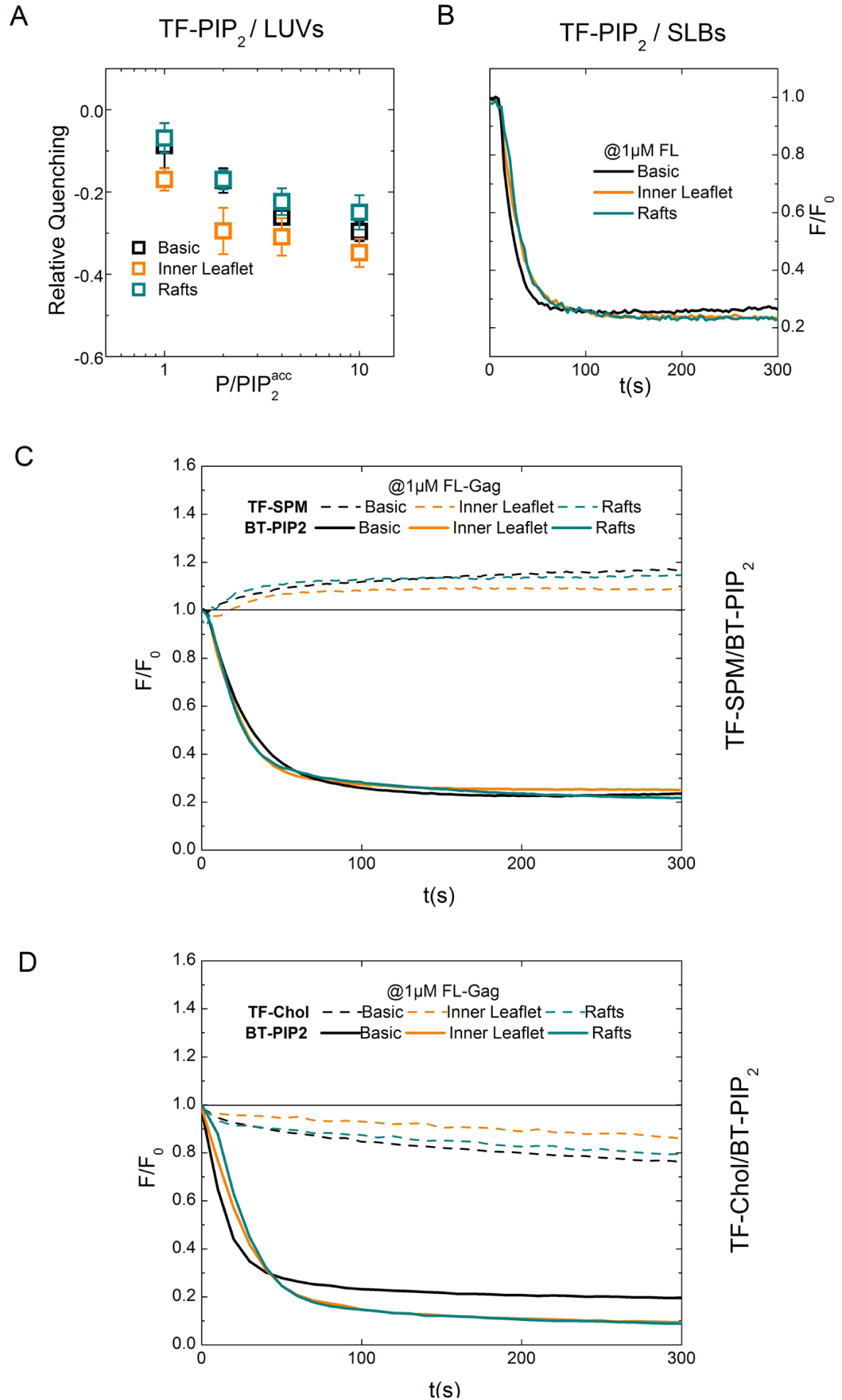
Gag PIP_2_ and Chol nanoclustering in complex membrane models. **A & B** TF-PIP_2_ fluorescence changes in lipid membranes of different composition (basic, Inner Leaflet, Raft) using LUVs (**A**: relative quenching of at different 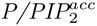 ratio. mean±s.d., n≥3) or SLBs (**B**: typical time course of TF-PIP_2_ fluorescence after addition of 1 µM FL-Gag.) **C**: Simultaneous fluorescence time course obtained on “inner leaflet” SLBs for BT-PIP_2_ (in red) and in green TF-SPM (left panel) or TF-Chol (right panel). **D**: Simultaneous fluorescence time course obtained on “raft” SLBs for BT-PIP_2_ (in red) and in green TF-SPM (left panel) or TF-Chol (right panel).

These results show that, as in the “basic” composition, in complex lipid mixtures Gag self-assembly induced PIP_2_ nanoclusters are enriched in Chol but not in SPM. This clearly suggest that Gag is sorting these lipids during self-assembly independently of the surrounding lipids chemical nature and acts as the driving force for lipid reorganization during HIV-1 assembly.

### Discussion

Although the MA domain is primary responsible for HIV-1 Gag binding to the PM, the ten times higher apparent affinity for membrane models observed here in the case of FL-Gag, P39 and WM confirms that the NC domains and CA-CA self-assembly of Gag are involved in membrane binding efficiency. Indeed, the NC domain of Gag alone has recently been described to bind to PIP_2_ containing lipid membranes with an apparent *K_p_* as high as 7 *µM* [35]. It has also already been shown that the driving force for membrane association stems largely from ionic interactions between multimerized Gag and negatively charged phospholipids [14].

Using micrometer scale phase separated GUVs, we have observed that Gag was mainly partitioning in L*_d_* phases, i.e., more likely outside of lipid “rafts” (namely Chol and SPM enriched) domains. The lack of myristate in the different Gag variants tested here could explain the Gag L_*d*_ phase localization [37]. Nevertheless, our results are consistent with the L_*d*_ phase GUV localization of multimerizable myr(+)MA protein observed by Keller *et al.* [34].

However, it is known that cholesterol is crucial for virus infectivity [9,26] and that Gag can sense cholesterol [4,16]. It was recently described that PIP_2_ could form clusters in the presence of cholesterol alone [33]. Here we show that PIP_2_ and cholesterol are laterally redistributed upon Gag self-assembly on membranes and participate to the formation of cholesterol/PIP_2_/Gag enriched nanodomains. Importantly, we also show that sphingomyelin is not sorted during HIV-1 Gag self-assembly and is excluded from the Gag/PIP_2_/Chol nanoclusters. Taken together, these results suggest that binding and self-assembly of Gag protein does not occur in pre-existing lipid domains (such as “rafts”) but, on the opposite that this self-assembly is more likely to induce lipid nanodomains. It is known that plasma membrane lipid organization is not only driven by lipid-lipid interactions. For example, cytoplasmic proteins such as ezrin [12] [1], syntaxin-1 [31] have also been described to induce or interact with PIP_2_ nanoclusters. As a general mechanism, proteins with basic interfaces can recruit acidic lipids that, in turn, can facilitate recruitment and clustering of these proteins into nanodomains [39,51]. A similar cooperative mechanism could also happen when HIV-1 Gag binds to PIP_2_ during viral assembly at the plasma membrane.

Interestingly, mesoscale organized structures such as subcortical actin cytoskeleton has also been shown to play an important role in the lateral organization of not only transmembrane proteins, but inner leaflet plasma membrane lipids such as PIP_2_ [23] and more interestingly outer leaflet components such as GPi-anchored proteins [24]. Moreover, subcortical actin cytoskeleton has also been shown to play a role in HIV-1 Gag assembly at the PM of Jurkat T-cells [50].

Since HIV-1 Gag has been often found into detergent resistant membranes (DRM) [44] pre-existing outer leaflet rafts could be trapped by these nascent PI(4,5)P2-Gag nanodomains through transbilayer coupling, as we already proposed [36] or as it has been recently shown for lipid domains [6] and for outer leaflet GPi anchored proteins [47]. Moreover, a recent study of the SPM dynamic in the cellular PM revealed that instead of clearly partitioning into nanodomains, SPM was mainly transiently trapped [30]. In the case of the ganglioside GM1, another lipid described as partitioning into L*_o_* phases, its transient trapping was shown to depend on molecular pinning and interleaflet coupling between lipid tail domains [49]. This suggest that Gag could not only generates his own lipid nanodomains at the inner leaflet of the cellular PM but also induce the formation of domains in the outer leaflet of the cellular PM by transiently trapping SPM or other components.

Finally, we also observed that the matrix domain of Gag is not able to induce this PIP_2_ clusterization, suggesting that, after maturation and particle release, the inner leaflet lipids of the virus envelope might be free to diffuse again.

## Conclusion

We have shown using simple and cell mimicking lipid composition model membranes that Gag self-assembly is inducing nanoclusters enriched in PIP_2_ and cholesterol instead of partitioning to pre-existing ones. This lipid nanoclusterization does not require sphingomyelin and mainly occurs out of the L*_o_* phases in GUVs. Further, the different lipid composition tested here does not strikingly affect the capacity of Gag to induce these lipid nanoclusters suggesting that Gag is able to sort his own lipids independently of the surrounding lipids.

## Methods

All experiments were performed at room temperature (RT). Experimental buffer was Hepes 10 mM, pH=7.4, KCl 150 mM, EDTA 2 mM except for QCM-D binding experiments. All images were acquired with a Zeiss LSM 780 microscope (Carl Zeiss, Inc.) using a 63x NA 1.4 oil objective and quantified using Image J software (NIH, MD, USA). Detailed materials, methods for model membranes (LUV, GUV, SLB) preparation and protein purification are in SI.

### Binding experiments

*K_p_* were determined on basic lipid composition (EPC 68%, BPS 30% & PI(4,5)P_2_ 2% for LUVs or POPC 68%, POPS 30% & PI(4,5)P_2_ 2% for SLBs). Methods used were either cosedimentation assays for LUVs or QCM-D experiments on SLBs. Co-sedimentation assays were made at 1 *µM* protein concentration with varying concentrations of total accessible lipids from 0.07 to 2250 *µM* in 100 *µl* of the same buffer that the one used in quenching experiments, according to the protocol in [25]. After 15 min of incubation at room temperature, samples were centrifuged at 220,000 g for 1 h at 4°C using a Beckman Coulter’s TLA 100 rotor. The top 80 *µl* was considered as supernatant (S) and the remaining 20 *µl* diluted with 60 *µl* of working buffer as pellet (P). Pellet and supernatant were analyzed on a 10% SDS-PAGE and stained using coomassie blue. Quantifications were made using Image J software (National Institutes of Health, MD, USA). SLBs were prepared with 0.1 mg.mL^−1^ liposomes flowing at 10 *µ*L.min^−1^ for 10-20 min on a UV-treated SiO_2_ surface of Q-sensor fixed in a Q-Sense Flow module, QFM 401 Biolin Scientific, Sweden). Stable SLBs were rinsed with citrate buffer (NaCitrate 10 mM, 100 mM NaCl, and 0.5 mM EGTA, pH 4.6) and then with injection buffer (5 mM Tris & 100 mM NaCl pH 7.4). At equilibrium, 200 *µ*L of increasing protein concentration was successively injected into the flow chamber followed by rinsing steps in between. The same was repeated with increasing protein concentrations until saturation. Sensorgrams were normalized to third harmonic in case of varying harmonic curves. *∆F* (plateau values) were used to measure the lipid SLB surface fraction of protein bound. The fraction of protein bound is related to an apparent association constant *K* (i.e, the reciprocal of the apparent partitioning constant, *K_p_*) following the equation:

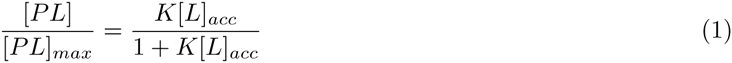

where the percentage of protein bound is [*PL*] = *I*_*Pellet*_/(*I_P_* + *I_S_*) in the case of the co-sedimentation experiments (*I_P_* & *I_S_* experimental intensities of pellet and supernatant and *I_Pellet_* = *I_P_* – 0.2.*I_S_*, for dilution compensation). In the case of QCM-d experiments, apparent *K_p_* values were obtained using Eq.1 formulated as a function of [*P*] instead of [*L*].

### Image analysis of GUVs

GUVs were deposited on coverslips coated with casein (2*mg.ml*^−1^) fitted in an Attofluor cell chamber (Thermo Fisher Scientific, Inc.). Labeled FL-Gag was added to the buffer. Spectral images were acquired at constant excitation intensity at 488 nm, 561 nm and 633 nm using an emission spectral range between 499 and 690 nm with a 8 nm resolution. Linear unmixing of these images was done to avoid potential bleed-through due to fluorophore emission overlapping. To establish the spatial auto-correlation of the fluorescence intensity of TF-PIP_2_ or A594-FL-Gag on basic composition GUVs, the change in intensity was plotted along the GUV using the following integration:

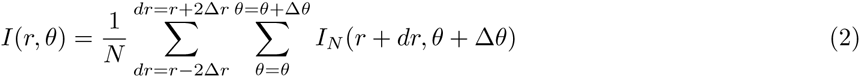

Intensity was the plotted with *r.sinθ* as the length unit. The obtained curved was autocorrelated using either the autocorr or the xcorr function of Matlab R2015 (Mathworks^®^).

Intensity partition of each label in the case of raft GUVs was determined using the following equations:

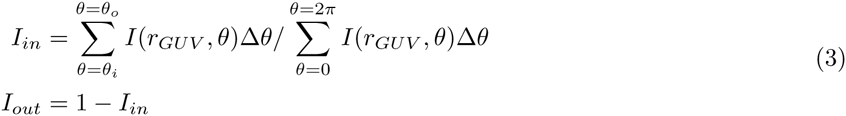

*θ_i_* and *θ_o_* were determined with the help of Alexa647-CtxB intensity circular profile.

### Fluorescence quenching measurements

For LUV experiments, fluorescence was monitored using a spectrometer (Photon Technologies International, Inc.) with *λ_exc_ =* 485 ± 2*nm* and *λ_em_* = 520 ± 10*nm*. Excitation lamp intensity was calibrated using the Raman spectrum of pure water and its fluctuations corrected every second. For every interaction assay on LUVs, the obtained intensity curve was corrected for both bleaching and dilution effect and then normalized to the mean intensity before injection. In the case of SLBs, images were acquired using the Zeiss definite focus system to avoid any z-drift with a 2-photons excitation (930 nm) every 5s in order to strongly reduce photobleaching. TF-lipids fluorescence was acquired at 520 ± 30 nm and BT-PIP_2_ at 600 ± 40. Mean intensity of each image was normalized to the intensity before injection.

### Self-assembly assay on LUVs

100 *µ*L of the basic LUVs was incubated at room temperature for 15 min with either 0.9 *µ*M or 4.5 *µ*M of FL-Gag or WM. After centrifugation at 10,000 g for 5 min at 4°C, 20 *µ*L of the supernatant was loaded on a 10% native-PAGE and proceed for immunoblot as in [25]. FL-Gag and WM were detected by a primary anti-capsid antibody (HIV-1 p24 NIH AIDS Reagents) followed by secondary antibody goat anti-mouse HRP conjugated. Membrane was revealed by Femto substrate (Thermo-scientific) and imaged by a G:Box (Syngene).

### Fluorescence Recovery After Photobleaching

The image sequence was acquired at 20 Hz using the 488 nm line of an Ar^+^ laser at a very low power to avoid photobleaching. After 2.5 s, 3 regions of interests (ROI), of 1*µ*m radius each, were rapidly photobleached (t*<*60 ms) at maximal laser power. Fluorescence recovery was monitored for 15s. The recovery curves were obtained as in [19]. In order to correctly estimate *F_0_* (the fluorescence intensity immediately after the end of the bleach) and *F*_15*s*_, the curves were fitted as in [19]. The fractional recovery (*FR*), is defined as:

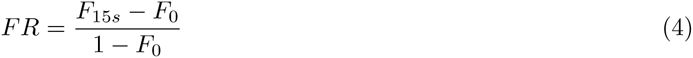

## Acknowledgements

We are thankful to Dr. Olivier Coux for his help with protein purification and Dr. Michel Franco for the gift of PH-EFA6 plasmid. We thank the Montpellier RIO Imaging (MRI) microscopy facilities. CF and DM are part of the CNRS GDR MIV consortium.

## Author contributions statement

DM and CF developed the concept of the study. NY, QL, HT, CF conducted the experiments. NY, QL and DM and CF conducted data interpretation. CF, NY and DM drafted the manuscript. JM, CP contributed to data interpretation and manuscript drafting.

## Supporting Information

Supporting information is provided in an separate file that contains 3 figures and a table.

